# Cortical processing of reference in language revealed by computational models

**DOI:** 10.1101/2020.11.24.396598

**Authors:** Jixing Li, Shaonan Wang, Wen-Ming Luh, Liina Pylkkänen, Yiming Yang, John Hale

## Abstract

Human language processing involves not only combining word meanings in accordance with semantic and syntactic constraints, but also figuring out who and what is being referred to. Here we present a first study towards a mechanistic understanding of the neural basis for referential processing. Using both functional MRI and magnetoencephalography (MEG), we identified a consistent increase of activity in a network spanning the anterior and posterior left middle temporal gyrus and the angular gyrus for pronoun processing during naturalistic listening for both English and Chinese speakers. We then adopted a “reverse-engineering” approach to examine the cognitive processes underlying pronoun resolution. We evaluated the neural fit of three symbolic models that each formalizes a different strand of explanation for pronoun resolution in the cognitive and linguistic literature, as well as two deep neural network models with an LSTM or a Transformer architecture. Our results favor the memory-based symbolic model, suggesting a domain-general mechanism of pronoun resolution that resembles memory retrieval.

## 1 Introduction

Word meaning involves not only the general, dictionary-like meaning (sense) but also the specific entity it denotes (reference). The classic example used by Frege (1892) is “the morning star”: “the morning star” and “the evening star” differ in their sense, but they have the same reference: the Venus. Reference allows us to relate our ideas of things to entities in the world. One example of this is using pronouns such as “she” and “it” to refer to a person and an object in the context. Pronouns are ubiquitous in human language, yet despite decades of research on pronoun resolution in linguistics and cognitive science, the neural basis for pronoun processing remains largely uncharacterised, and it is absent from recent neurobiological models of language comprehension (e.g., Friederici, 2011; Hagoort & Indefrey, 2014). Prior neuroimaging studies on referential processing have reported a number of regions, including the medial parietal lobe (van Berkum et al., 2007; Brodbeck & Pylkkänen, 2017; Brodbeck et al., 2016), the lateral parietal region (van Berkum et al., 2007), the inferior frontal region (Hammer et al., 2007; Matchin et al., 2014; Miceli et al., 1991) and the temporal regions (Hammer et al., 2007, 2011; Miceli et al., 1991), yet they involve different experimental manipulations such as contrasting referentially ambiguous pronouns with non-ambiguous pronouns (van Berkum et al., 2007), contrasting gender-incongruent with gender-congruent pronouns (Hammer et al., 2007). It is therefore unclear whether these manipulations tapped the same cognitive processes.

As for the cognitive processes relevant for pronoun resolution, various accounts have been proposed in linguistics and cognitive science. The classic Binding Theory (Chomsky, 1981) in formal linguistics suggests that pronoun resolution is constrained by syntactic constraints, such that pronouns cannot be coindexed with antecedents in the same clause. For example, in “Mary loves her”, the pronoun “her” cannot refer back to the “Mary”. The Centering theory (Grosz et al., 1995) in pycholinguisitcs emphasizes the role of discourse coherence when interpreting pronouns and claims that pronouns refer to the most prominent entity in the discourse context. The memory-based account (Nieuwland & van Berkum, 2006) suggests that pronoun resolution involves maintaining different references in the working memory, hence is influenced by memory resources. These accounts each emphasizes a different aspect of how we link a pronoun to an antecedent expression during language comprehension, yet it is unknown which account best accounts for the brain mechanism of pronoun resolution.

Here we conducted two analyses to examine both the neural correlates and the cognitive processes for pronoun resolution (see Figure 1). We recorded the blood-oxygen-level-dependent (BOLD) signals while both English and Chinese participants listened to the audiobook of “The Little Prince” for about 100 minutes in the fMRI scanner. This naturalistic setting allows us to examine pronoun processing without invoking ungrammatical constructions. The cross-linguistic comparisons between English and Chinese also provide insights on whether linguistic typology would influence the strategies for pronoun resolution, as Chinese pronouns do not distinguish gender in its spoken form (“she”, “he” and “it” are all pronounced as “ta” in Chinese). To further examine the temporal dynamics of pronoun processing, we collected a magnetoencephalography (MEG) dataset while English speakers listened to a 12-minute audio excerpt from the YouTube channel “SciShow Kids”. We conducted a general linear model (GLM) analysis to localize the fMRI BOLD signals and the MEG source estimates time-locked to each third person pronoun in the narratives. We identified a consistent increase of activity in the anterior and posterior left middle temporal gyrus (MTG) and the angular gyrus (AG) across languages and imaging methods, starting from around 150 ms to 250 ms after the onset of the pronouns.

**Figure 1:**
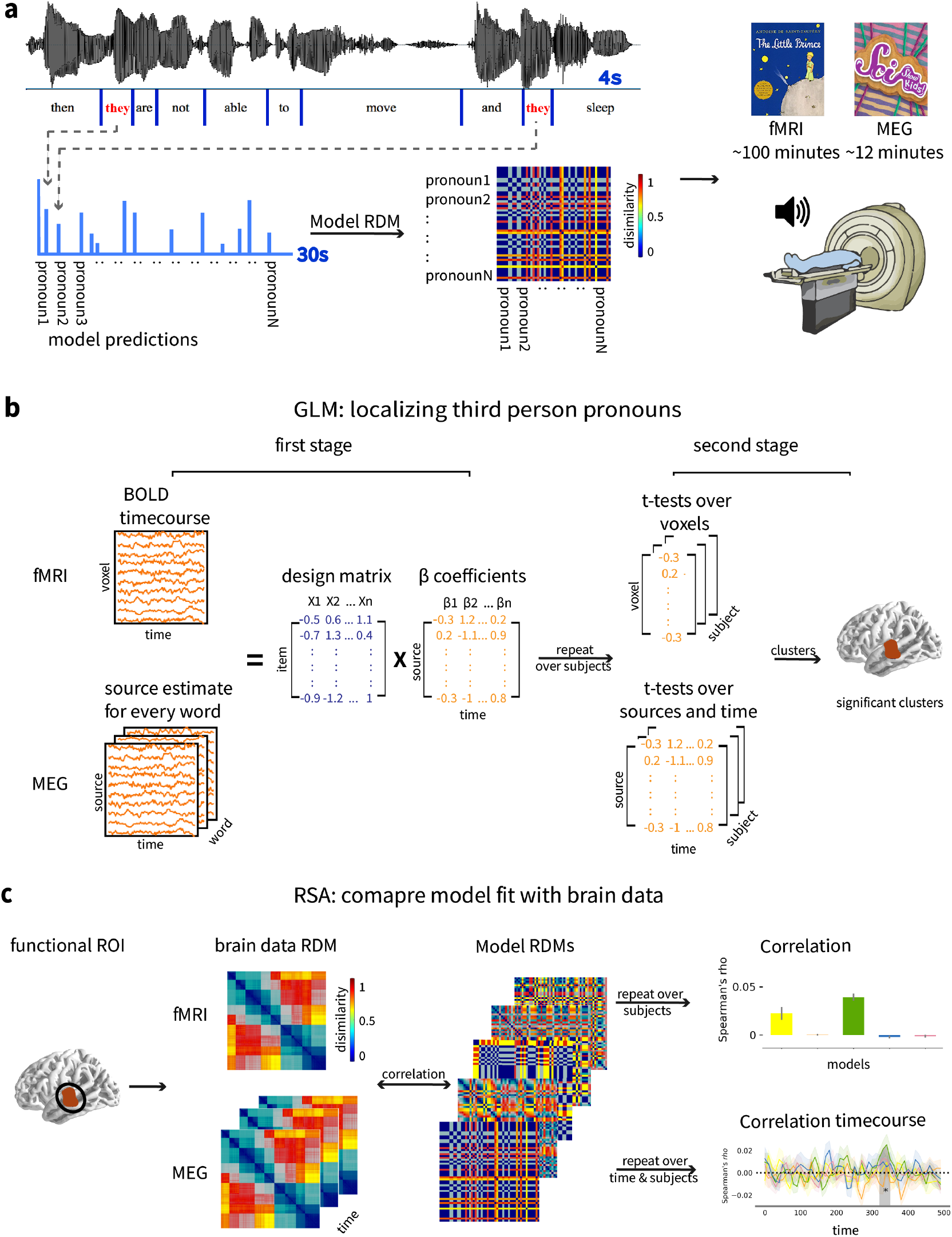
Schematic illustration of the analysis pipeline. **a** Participants listened to the narrative in the fMRI/MEG scanner. Model predictions for all the third person pronouns in the narratives. **b** Localizing third person pronoun processing using GLM. **c** Comparing model relatedness to fMRI/MEG activity pattern within the fROIs derived from the GLM analyses using RSA.

To understand pronoun processing at the algorithmic level, we adopted the integrative “reverse-engineering” approach that links computational models with brain functions. Computational models provide a viable way of specifying complex and detailed theories of the underlying cognitive process. Consequently, they make quantitative predictions that can be rigorously tested against human brain activity. This “reverse-engineering” approach has been successfully applied to understand the neural mechanisms of primate and human vision (e.g., Bao et al., 2020; Cadena et al., 2019; Kietzmann et al., 2019), as well as the higher-level cognition of language (e.g., A. Anderson et al., n.d.; J. Brennan et al., 2012, 2016; Jackson et al., 2021; Hale et al., 2018; Martin & Doumas, 2019; Toneva & Wehbe, 2019; Toneva et al., 2021; Schrimpf et al., 2021; Wehbe et al., 2014). Here we compared the fit of five computational models for pronoun processing with the fMRI and MEG data within the LMTG functional regions of interest (fROIs) using representational similarity analysis (Kriegeskorte et al., 2008). Three of the models are knowledge-based symbolic models that formalize the Binding Theory (Hobbs, 1977), the Centering Theory (S. Brennan et al., 1987) and the memory account for pronoun resolution (van Rij et al., 2013), while the other two models are data-driven deep neural networks with either a long short-term memory (LSTM) architecture (ELMo; Lee et al., 2017; Lee et al.,2018) or a Transformer architecture (BERT; Joshi et al., 2019). Although ELMo and BERT are not intended to be cogntive models of lanaguage processing, they have shown high performance in Natural Language Processing (NLP) and hold great promise for simulating the brain’s language system (Toneva et al., 2021; Schrimpf et al., 2021). All the models can adequately predict the the correct antecedent for the third person pronouns in the English and Chinese fMRI stimuli and the English MEG stimuli (see Section 2.4). However, when compared with brain data, all our results favored the memory-based ACT-R model, suggesting a domain-general mechanism for pronoun resolution that resembles memory retrieval.

## 2 Materials and methods

### 2.1 Participants

The English fMRI data were taken from a published study (Bhattasali et al., 2018), collected at the same time with the Chinese data. Participants were 49 young adults (30 females, mean age = 21.3, SD=3.6) with no history of psychiatric, neurological or other medical illness that might compromise cognitive functions. They self-identified as native English speakers, and strictly qualified as right-handed on the Edinburgh handedness inventory (Oldfield, 1971). All participants were paid, and gave written informed consent prior to participation, in accordance with the IRB guidelines of Cornell University.

Chinese participants of the fMRI study were 35 young adults (15 females, mean age=19.3, SD=1.6) with normal hearing and no history of psychiatric, neurological or other medical illness that might compromise cognitive functions. They self-identified as native Chinese speakers, and strictly qualified as right-handed on the Edinburgh handedness inventory (Oldfield, 1971). All participants were paid, and gave written informed consent prior to participation, in accordance with the IRB guidelines of Jiangsu Normal University.

Participants for the MEG study were 13 young adults (7 females, mean age=19.9, SD=1.3) with normal hearing and no history of psychiatric, neurological or other medical illness that might compromise cognitive functions. They self-identified as native English speakers, and strictly qualified as right-handed on the Edinburgh handedness inventory. All participants were paid, and gave their written informed consent prior to participation, in accordance with New York University Abu Dhabi IRB guidelines.

### 2.2 Stimuli and annotation

The English fMRI stimulus is a 94-minute audiobook of Antoine de Saint-Exupéry’s *The Little Prince*, translated by David Wilkinson and read by Nadine Eckert-Boulet. The Chinese fMRI stimulus is a 99-minute Chinese translation of *The Little Prince* (“小王子网站”, 2020), read by a professional female Chinese broadcaster hired by the experimenter. The MEG stimulus is a 12-minute audio excerpt taken from the YouTube channel “SciShow Kids”. It consists of 4 short audios that introducing scientific fun facts to kids. Word boundaries in the audios were identified and aligned to the transcripts using the Forced Alignment and Vowel Extraction (FAVE) and were manually checked by native English and Chinese speakers.

All entities (or mentions; e.g., nouns, pronouns, proper names, etc.) in the audio texts were first identified using the Stanford Named Entity Recognizer (NER) (Finkel et al., 2005). They were then manually checked and linked with their coreferential mentions using the annotation tool brat (Stenetorp et al., 2012). Supplemental Figure 1 demonstrates sample annotations for the three texts. We identified 4882 mentions in the English fMRI stimulus, 4732 mentions in the Chinese fMRI stimulus, and 701 mentions in the MEG stimulus. We then selected only the third person pronouns (e.g., he, him, she, her, it, they, them) from the mentions as our focus of analyses as the Hobbs and Centering models for pronoun resolution are mainly concerned with third person pronouns. In addition, the cognitive processes may differ for first, second and third person pronoun processing (Ariel, 1990). We removed possessives, reflexives, cleft and extraposition “it”, pleonastic “it”and pronouns with sentential antecedents. The final English fMRI stimulus contains 647 third person pronouns, the Chinese fMRI stimulus contains 524 third person pronouns, and the MEG stimulus contains 51 third person pronouns (see Supplemental Figure 2).

### 2.3 Computational models for pronoun resolution

To gain a mechanistic understanding of pronoun processing in the brain, we compared both BOLD responses and MEG source estimates for pronoun processing with five computational models that specifies the detailed steps for pronoun processing. We selected three symbolic models that each formalizes a different strand of explanation for pronoun resolution that has figured in the cognitive and linguistic literature: the syntax-based Hobbs model (Hobbs, 1977), the discourse-based Centering model (S. Brennan et al., 1987) and the memory-based ACT-R model (van Rij et al., 2013). We also included two deep neural network models with architectures that have been shown to simulate the brain’s language system (Toneva et al., 2021; Schrimpf et al., 2021): the ELMo-Coref model (Lee et al., 2017, 2018) uses bidirectional LSTMs to capture the contextual information, and the BERT-Coref model (Joshi et al., 2019) uses a Transformer architcuture (Devlin et al., 2019). See Section 2.3.1–5 for a detailed description of the models. Figure 2 shows an illustration of the four models applied to an example English sentence from “The Little Prince”.

**Figure 2:**
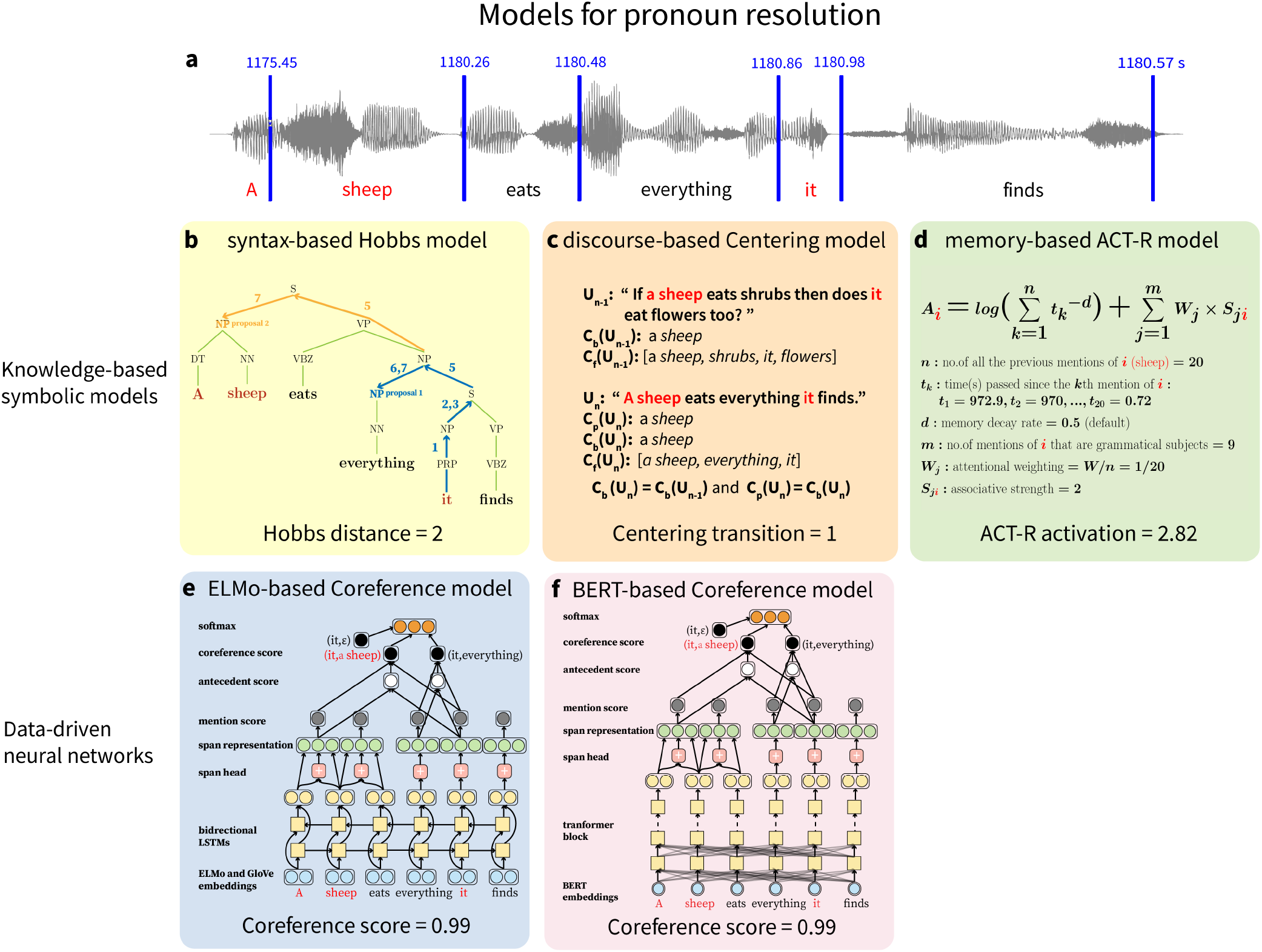
Illustration of the Hobbs (Hobbs, 1977), Centering (S. Brennan et al., 1987), ACT-R (van Rij et al., 2013), ELMo-Coref (Lee et al., 2017, 2018) and BERT-Coref (Joshi et al., 2019) models using an example sentence in the English *The Little Prince*. **a** Waveform of the example sentence from the English audiobook. The blue numbers indicate the offset time in seconds for each word in the whole audiobook. **b** Hobbs model applied to the English example sentence. Blue arrows indicate the steps performed to get to the first proposed antecedent, and the orange arrows indicate the steps performed to get the second proposed antecedent. Hobbs distance is the number of proposals till the correct antecedent. **c** Centering model applied to the English example sentence. *C_b_* equals *C*_*b*–1_ and *C_p_* equals *C_b_*, so the transition type is Continuing and the transition ordering is 1. **d** The formula for calculating the activation level for the antecedent “a sheep”. **e** The architecture of the ELMo-Coref model (Lee et al., 2017, 2018). The coreference score is the softmax of the final layer. **f** The architecture of the BERT-Coref model (Joshi et al., 2019). The coreference score is the softmax of the final layer.

#### 2.3.1 The Hobbs model

The Hobbs model for pronoun resolution (Hobbs, 1977) depends on a syntactic parser plus a morphological gender and number checker. The input to the Hobbs model includes the target pronoun and the parsed trees for the current and previous sentences. The model searches for a gender and number matching antecedent by traversing the trees in a left-to-right, breadth-first order, that is, it starts at the tree root and explores the neighboring nodes at the present depth prior to moving on to the nodes at the next depth level. If no candidate antecedent is found in the current tree, the algorithm searches on the preceding sentence in the same order. When applied to the Chinese text, the Hobbs model no longer contains a gender and number agreement checker because pronouns in spoken Chinese do not distinguish gender and Chinese NPs usually do not mark plurals. The steps of the Hobbs algorithm are as follows:

1. Begin at the NP node immediately dominating the pronoun.
2. Go up the tree to the first NP or S node encountered. Call this node X, and call the path used to reach it p.
3. Traverse all branches below node X to the left of path p in a left-to-right, breadth-first fashion. Propose as the antecedent any NP node that is encountered which has an NP or S node between it and X.
4. If node X is the highest S node in the sentence, traverse the parsed trees of previous sentences in the text in order of recency, the most recent first; each tree is traversed in a left-to-right, breadth-first manner, and when an NP node is encountered, it is proposed as antecedent. If X is not the highest S node in the sentence, continue to step 5.
5. From node X, go up the tree to the first NP or S node encountered. Call this new node X, and call the path traversed to reach it p.
6. If X is an NP node and if the path p to X did not pass through the *N* node that X immediately dominates, propose X as the antecedent.
7. Traverse all branches below node X to the left of path p in a left-to-right, breadth-first manner. Propose any NP node encountered as the antecedent.
8. If X is an S node, traverse all branches of node X to the right of path p in a left-to-right, breadth-first manner, but do not go below any NP or S node encountered. Propose any NP node encountered as the antecedent.
9. Go to step 4.

The Hobbs model incorporates the locality constraints of the Binding Theory (Chomsky, 1981) as it always searches for the antecedent in the left of the noun phrase (NP) and does not go below any NP or S(entence) Node on the tree. We used the “Hobbs distance” (Ge et al., 1998) metric to represent the processing complexity of the pronouns derived by the Hobbs model. Hobbs distance refers to the number of proposals that the Hobbs algorithm has to skip, starting backward from the pronoun, before the correct antecedent NP is found. Figure 2c,d illustrate the Hobbs model for one example sentence in English. For the English sentence, the model first proposes the noun phrase (NP) “everything” as the antecedent of “it”, which is incorrect, so the Hobbs distance is 2.

#### 2.3.2 The Centering model

The Centering model for pronoun resolution (S. Brennan et al., 1987) (also known as the BFP algorithm) formalizes the Centering Theory (Grosz et al., 1995). In the Centering framework, entities that link an utterance to other utterances are referred to as “centers”. Centers of an utterance are ranked according to their relative prominence, which is mainly determined by the centers’ grammatical roles. In particular, subjects of a sentence rank higher than objects, and objects rank higher than other grammatical roles. Each utterance (*U_n_*) has a set of forward-looking centers (*C_f_*) and a single backward-looking center (*C_b_*). *C_f_* (*U_n_*) contains all the entities in *U_n_* and *C_b_* is the highest-ranked entity among the entities in the previous utterance (*C_f_* (*U*_*n*–1_)). The transition relations between the forward- and backward-looking centers in an adjacent pair of sentences are classified into three types: Continuing, Retaining and Shifting. In the Continuing transition, propositions of the current entity are maintained, that is to say, *C_b_*(*U_n_*) is the same entity as the backward-looking center of the previous utterance (*C_b_*(*U_n_*) = *C_b_*(*U*_*n*–1_)), and *C_b_*(*U_n_*) is also the preferred center of the current utterance (*C_p_*(*U_n_*)), i.e., the highest-ranked entity in *C_f_* (*U_n_*) (*C_b_*(*U_n_*) = *C_p_*(*U_n_*)). In the Retaining transition, a related entity is introduced to the context, thus *C_b_*(*U_n_*) is the same as *C_b_*(*U*_*n*–1_), but it is not the highest-ranked entity in *C_f_* (*U_n_*) (*C_b_*(*U_n_*) = *C_b_*(*U*_*n*–1_) and *C_b_*(*U_n_*) = *C_p_*). In the Shifting transition, a new entity becomes the center of the discourse, therefore, *C_b_* (*U_n_*) is not the same entity as *C_b_* (*U*_*n*–1_) (*C_b_*(*U_n_*) = *C_b_*(*U*_*M*–1_)). For a discourse segment to be coherent, Continuing is preferred over Retaining, which is preferred over Shifting. Frequent Shifting leads to a lack of discourse coherence and substantially affects the processing demands made upon a hearer during discourse comprehension.

The Centering Theory claims that pronominalization serves to increase discourse coherence and eases the hearer’s processing difficulty of inference. Based on this assumption, the Centering model (S. Brennan et al., 1987) tracks the relation between the forward- and backward-looking centers in adjacent pairs of sentences and finds the antecedent-pronoun pair that has the highest-ranked transition types. The model further divides Shifting into Shifting where *C_b_*(*U_n_*) = *C_b_*(*U*_*n*–1_) and *C_b_*(*U_n_*) = *C_p_*, and Shifting-1, where *C_b_*(*U_n_*) = *C_b_*(*U*_*n*–1_) and *C_b_*(*U_n_*) = *C_p_*. The coherence ordering of the transition types is Continuing > Retaining > Shifting > Shifting. The algorithm consists of three basic steps. In the first step, it constructs all possible *C_b_–C_f_* pairs for the pronoun in the current utterance (*U_n_*). These pairs are called “anchors” and they represent all the coreferential relationships available for this utterance. Next, the model filters the anchors based on the Centering rule that *C_b_* must be pronominalized if any *C_f_* is pronominalized. Finally, the algorithm ranks the remaining pairs by the ordering of the transitions and select the pair that has the preferred transition types. The rank of the entities in *C_f_* is determined by their grammatical roles, which is parsed using the Stanford dependency parser for English (de Marneffe et al., 2006) and Chinese (Chang et al., 2009).

We used the rank of the transition types for each correct antecedent-pronoun pair generated by the Centering model to indicate the processing difficulty of the pronouns. Figure 2c illustrates how the model classifies the transition type for the example discourse segment in “The Little Prince”. The current utterance (*U_n_*) “A sheep eats everything it finds” has a set of *C_f_* (*U_n_*): (“a sheep”, “everything”, “it”); the preferred center (*C_p_*(*U_n_*)) is “a sheep” as it is the subject of the sentence. The backward-looking center of the current (*C_b_*(*U_n_*)) and the previous utterance (*C_b_*(*U*_*n*–1_)) is also “a sheep” as it is the subject of the *U*_*n*–1_ (highest-ranked entity in *C_f_* (*U*_*n*–1_)). Since *C_b_*(*U_n_*) = *C_b_*(*U*_*n*–1_) and *C_b_*(*U_n_*) = *C_p_*(*U_n_*), the transition type is CONTINUING and the rank is 1.

#### 2.3.3 The ACT-R model

The ACT-R model for pronoun resolution (van Rij et al., 2013) uses the same primitives of the memory module in the cognitive architecture of ACT-R (J. R. Anderson, 2007). The formula for the activation level of antecedent *i* of a pronoun is as follows:

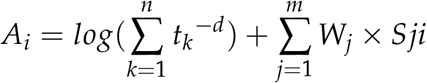

The first part of the equation 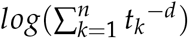 computes the inherent strength, or the base-level activation of entity *i*, which reflects the past usage of entity *i* in the text. *t_k_* is the time passed since the *k*th mention of *i*, and each mention decays over time as a negative power function *t_k_^−d^*. The parameter d is set to 0.5 as the default value in ACT-R based on a range of experiments to model human performance in memory retrieval tasks (J. R. Anderson, 2007). Different mentions of the entity i are summed up to reflect the effect of practice. We calculated the base-level activation of each pronoun in English and Chinese based on the offset time of each pronoun and their previous mentions in the audio.

The second part of the equation 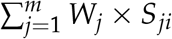 reflects the associative activation that entity i receives from the mentions of *i* that are the subjects of their sentences. *S_ji_* is the strength of association reflecting how much the presence of each subject mention *j* of entity *i* makes *i* more salient. The value of *S_ji_* is set to 2 in our implementation. *W_j_* is the attentional weighting which equals to *W/n*, where *n* is the number of all the previous mentions of *i*, as the total value of associative activation cannot be infinite. The attentional weight *W* is set to 1. Subjecthood of each mention in the English and Chinese texts was annotated using the Stanford dependency parser (de Marneffe et al., 2006; Chang et al., 2009).

The effects of frequency and recency are folded into the calculation of the base activation for antecedent *i*, such that the more frequent and recent the previous mentions, the higher the base activation. Conversely, if antecedent *i* has been mentioned only once, or if it was mentioned a long time ago, its activation level will be lower, and it will rank lower on the activation list for all the candidate antecedents. Subjecthood of the previous mentions of antecedent *i* gains an associative activation in addition to the base activation. Overall, the activation level of an entity in the discourse context is computed based on recency, frequency and the grammatical role of the entity. The entity that has the highest activation level is predicted to be the antecedent of the pronoun. Figure 2d shows how the ACT-R activation level for a pronoun in the English example sentence is calculated.

#### 2.3.4 The ELMo-Coref model

The ELMo-based coreference model (Lee et al., 2017, 2018) is an end-to-end coreference resolution system that predicts all clusters of coreferential mentions given a text document. The model considers all possible spans in a document *D* containing *T* words as potential mentions. The total number of possible text spans in *D* is *N* = *T*(*T* + 1)/2. The task for the model is to assign an antecedent *y_i_* for each span *i* with a start *START*(*i*) and an end index *END*(*i*) for 1 ≤ *i* ≤ *N*. The possible assignments of each *y_i_* is *Y*(*i*) = {*ϵ*, 1,…, *i* – 1}. The model then learns a conditional probability distribution over the possible antecedents using the pairwise score *s*(*i, y_i_*) for each antecedent-span pair:

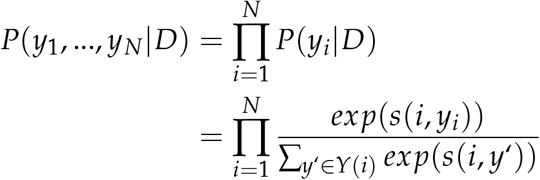

The architecture of the model can be divided into two parts. In the first part, the model encodes every word in its context using bidirectional LSTMs (Hochreiter & Schmidhuber, 1997):

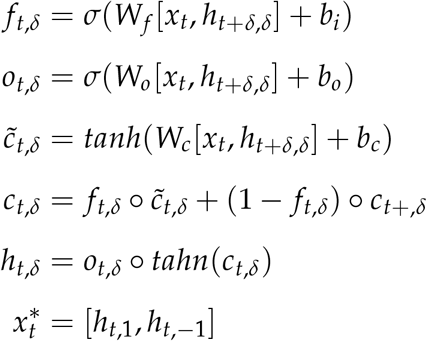

The model then assigns weights to the word vectors to represent the notion of syntactic head using an attention mechanism (Bahdanau et al., 2016):

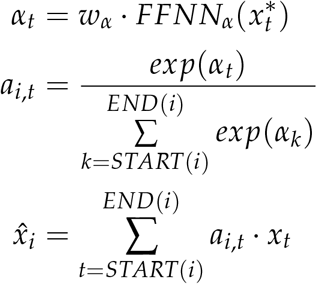

The context-dependent word embeddings and their weighted sum are concatenated to produce the span representations. In the second part, the model assigns a mention score to each span representation and an antecedent score to each antecedent-span pair via standard feed-forward neural networks. The antecedent scoring function incorporates a feature vector encoding speaker and genre information and the distance between the two spans. The mention score and the antecedent score are concatenated to produce the coreference score between the candidate antecedent and the mention span. During training, the model optimizes the marginal log-likelihood of all correct antecedents in the correct clustering *GOLD*(*i*) (see Figure 2e for the architecture of the model):

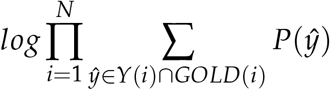

The input layer of the model consists of a fixed concatenation of the ELMo (Matthew et al., 2018) and GloVe embeddings (Pennington et al., 2014). The hidden layers in the LSTMs have 200 dimensions, and the two hidden layers in the feed-forward neural network have 150 dimensions. The speaker, genre, span distance and span width information are represented as 20-dimensional embeddings. To maintain computing efficiency, the maximal span width was set to 10, the maximal number of antecedent was set to 250 and the maximal number of sentences in the document was set to 50.

The model was trained on the English data from the CoNLL-2012 shared task (Pradhan et al., 2012) and achieved an average F1 score of 75.8. We took the pretrained English model (Lee et al., 2017, 2018) to generate the softmax of the coreference scores for all the pronouns in the English “The Little Prince” and the “SciShow Kids” texts. We then trained the model on the Chinese data from the CoNLL-2012 shared task (Pradhan et al., 2012). We used the pretrained ELMo embeddings for Chinese (Che et al., 2018) and the 300-dimensional word2vec embeddings (Li et al., 2018) trained on Baidu Encyclopedia. The model achieved an average F1 score of 63.1 for a single model on the test set of the Chinese data from the CoNLL-2012 shared task. We then took the trained Chinese model to generate the softmax of the coreference scores for all the pronouns in the Chinese “The Little Prince”.

#### 2.3.5 The BERT-Coref model

The BERT-based coreference model (Joshi et al., 2019) is an extension of the ELMo-based coreference model (Lee et al., 2017, 2018). It simply replaces the ELMo and GloVe embeddings and the LSTM encoders in the ELMo-Coref model with the BERT Transformers (Devlin et al., 2019). BERT’s model architecture is a multi-layer bidirectional Transformer encoder (Vaswani et al., 2017). The BERT-base model consists of 12 identical layers, and each layer contains a multi-head self-attention layer and a fully connected feed-forward network. The attention mechanism maps queries (Q) and a set of key-value (K,V) pairs to an output using a scaled dot-product function, where the queries and keys are vectors of dimension d_k_ and the values are vectors of dimension *d_v_*:

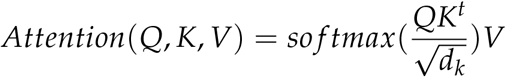

BERT-base employed 12 parallel attention layers, or heads, and the output from a single attention function are concatenated and feed to the feed-forward network layer:

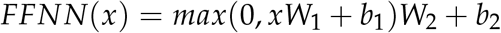

We used the pretrained BERT-base model for the English data (Joshi et al., 2019) to generate the softmax of the coreference scores for all the pronouns in the English “The Little Prince”. We then trained the same model on the Chinese data from the CoNLL-2012 shared task (Pradhan et al., 2012). The model achieved an average F1 score of 65.11 for a single model on the test set of the Chinese data from the CoNLL-2012 shared task. We took the trained Chinese model to generate the softmax of the coreference scores for all the pronouns in the Chinese “The Little Prince”.

### 2.4 Model comparisons

Since the features used by each model vary across pronouns in the narratives, the models predict different “processing difficulty” for each pronoun. For example, pronouns intervened by another candidate antecedent on a syntactic tree are hard for the Hobbs model; pronouns not referring to the most prominent entity in the discourse context are hard for the Centering model; pronouns referring to entities that are mentioned only a few times a while back are hard for the ACT-R model, and pronouns surrounded by nouns with similar word meanings are hard for the ELMo-Coref and the BERT-Coref model. To connect properties of the four models to the observed brain data, we define a “complexity metric” for each model to quantify how difficult it is for the model to find the correct antecedent. For the Hobbs model, we used the “Hobbs distance” metric (Ge et al., 1998), namely, the number of proposals that the Hobbs algorithm has to skip before the correct antecedent is found. For the Centering model, we used the rank of the transition type from the previous sentence to the current sentence containing the pronoun. For the ACT-R model, we used the negative of the activation level for the antecedent of each pronoun, and for the ELMo-Coref and the BERT-Coref models, we used the negative of coreference score of the antecedent for each pronoun. These complexity metrics allow us to estimate model-derived brain states for comparison against observed brain data.

Figure 3a shows the distribution of the standardized complexity metrics derived by the five models applied to all the third person pronouns in “The Little Prince” in English and Chinese and “SciShow Kids”. The distribution of the complexity metrics are very similar for the “The Little Prince” in English and Chinese, where the metrics are all right-skewed with more positive values of 2 or 3 z-scores. This suggest that texts contain more difficult pronouns for all the models. For the MEG stimuli, the complexity metrics are left-skewed with more negative values, suggesting more easier pronouns for the models. However, there are more pronouns with a complexity *z*-score of 1 than pronouns with mean complexity (*z*score=0), indicating more pronoun with medium difficulty levels in the MEG stimuli.

**Figure 3:**
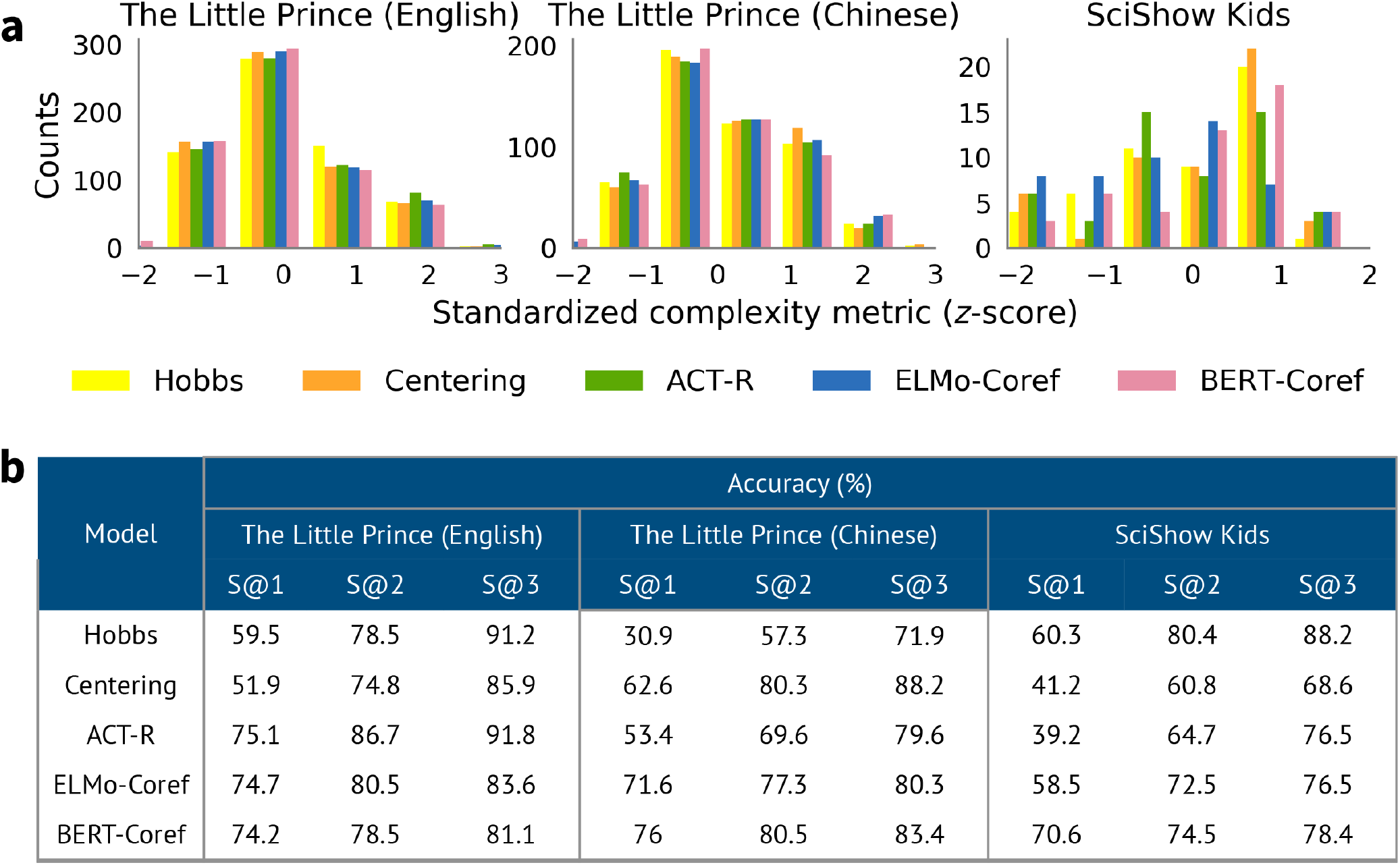
Comparison of the five models applied to the fMRI and MEG stimuli. **a** Distribution of the standardized model metrics. We took the negative of the ACT-R, ELMo-Coref and BERT-Coref metrics to indicate the “processing difficulty” of the pronouns, aligning with the Hobbs and the Centering metrics. All the metrics are z-scored. **b** The accuracy of the Hobbs, ACT-R, ELMo-Coref and BERT-Coref models based on SUCCESS@N (N=1,2,3) (Kolhatkar & Hirst, 2014), i.e., the proportion of the correct antecedent occurs within the model’s first three choices.

To ensure that the five models can indeed predict the correct antecedents, we calculated the accuracy of the five models applied to all the third person pronouns in the narratives. We allowed some degree of ambiguity in the reference and permitted the correct answer to rank within a model’s top 3 choices. This is the SUCCESS@N metric (Kolhatkar & Hirst, 2014) where the gold answer occurs within a system’s first N choices. All the models performed well with an accuracy well above or near 70% at SUCCESS@3 (see Figure 3b).

### 2.5 Experiment procedures

#### 2.5.1 fMRI experiments

After giving their informed consent, participants were familiarized with the MRI facility and assumed a supine position on the scanner. Auditory stimuli were delivered through MRI-safe, high-fidelity headphones (English: Confon HP-VS01, MR Confon, Magdeburg, Germany; Chinese: Ear Bud Headset, Resonance Technology, Inc, California, USA) inside the head coil. The headphones were secured against the plastic frame of the coil using foam blocks. An experimenter increased the sound volume stepwise until the participants could hear clearly. Both the English and Chinese stimuli were divided into 9 sections, and each lasted for about 10 minutes. Participants listened passively to the 9 sections and completed 4 quiz questions after each section (36 questions in total). These questions were used to confirm their comprehension and were viewed by the participants via a mirror attached to the head coil and they answered through a button box. The entire session, including preparation time and practice, lasted for around 2.5 hours.

#### 2.5.2 MEG experiment

After giving their informed consent, each participant’s head shape was digitized using a Polhemus dual source handheld FastSCAN laser scanner (Polhemus, VT, USA). Participants then completed the experiment while lying supine in a dimly lit, magnetically shielded room. MEG data were recorded continuously using a whole-head 208 channel axial gradiometer system (Kanazawa Institute of Technology, Kanazawa, Japan). Auditory stimuli were delivered through MEG-safe, high-fidelity earphones inside the head coil. The headphones were secured against the plastic frame of the coil using foam blocks. An experimenter increased the sound volume stepwise until the participants could hear clearly. The audio lasted for about 12 minutes. Participants listened passively to the audio and completed four picture-matching tasks. These tasks were used to confirm their comprehension and were completed by the participants outside the MEG scanner. The entire session lasted for around 30 minutes. All presentation scripts were written in PsychoPy2 (Peirce, 2007).

### 2.6 Data acquisition and preprocessing

#### 2.6.1 fMRI data

Both English and Chinese MRI images were acquired with a 3T MRI GE Discovery MR750 scanner with a 32-channel head coil. Anatomical scans were acquired using a T1-weighted volumetric Magnetization Prepared RApid Gradient-Echo (MP-RAGE) pulse sequence. Functional scans were acquired using a multi-echo planar imaging (ME-EPI) sequence with online reconstruction (TR=2000 ms; TEs=12.8, 27.5, 43 ms; FA=77°; matrix size=72 x 72; FOV=240.0 mm x 240.0 mm; 2 x image acceleration; 33 axial slices, voxel size=3.75 x 3.75 x 3.8 mm). All fMRI data were preprocessed using AFNI version 16 (Cox, 1996). The first 4 volumes in each run were excluded from analyses to allow for T1-equilibration effects. Multi-echo independent components analysis (ME-ICA) (Kundu et al., 2012) were used to denoise data for motion, physiology and scanner artifacts. Images were then spatially normalized to the standard space of the Montreal Neurological Institute (MNI) atlas, yielding a volumetric time series resampled at 2 mm cubic voxels.

#### 2.6.2 MEG data

MEG data were recorded continuously at a sampling rate of 1000 Hz with an online bandpass filter of 0.1-200 Hz. The raw data were first noise reduced via the Continuously Adjusted Least-Squares Method (Adachi et al., 2001) and low-pass filtered at 40 Hz. Independent component analysis (ICA) was then applied to remove artifacts such as eye blinks, heart beats, movements, and well-characterized external noise sources. The MEG data were then segmented into 500 ms epochs at the onset of each word in the stimulus. The epochs were baseline-corrected using the mean of the whole epoch. Epochs containing amplitudes greater than an absolute threshold of 2000 fT were automatically removed.

Cortically constrained minimum-norm estimates (Hämäläinen & Ilmoniemi, 1994) were computed for each epoch for each participant. To perform source localization, the location of the participant’s head was coregistered with respect to the sensor array in the MEG helmet using FreeSurfer’s (“Freesurfer”, 2020) “fsaverage” brain, which involved first rotation and translation and then scaling the average brain to match the size of the head scan. A source space of 2562 source points per hemisphere was generated on the cortical surface for each participant. The Boundary Element Model (BEM) was employed to compute a forward solution, explaining the contribution of activity at each source to the magnetic flux at the sensors. Channel-noise covariance was estimated based on the whole epoch. The inverse solution was computed from the forward solution and all the epochs. To lift the restriction on the orientation of the dipoles, the inverse solution was computed with “free” orientation, meaning that the inverse operator places three orthogonal dipoles at each location defined by the source space. When computing the source estimate, only activities from the dipoles perpendicular to the cortex were included. The same inverse operator was applied to each single trial to yield the dynamic statistical parameter maps (dSPM) units using an SNR value of 3. All data preprocessing steps were performed using MNE-python (v.0.19.2) (Gramfort & et al., 2014).

### 2.7 Localizing brain regions for third person pronoun processing

For the fMRI data, we modeled the timecourse of each voxel’s BOLD signals for each of the nine sections by a binary third person pronoun regressor, time-locked to the offset of each third person pronoun in the audiobook. We included four control variables: the root mean square intensity (`RMS intensity`) for every 10 ms of each audio section, the f0 of each audio section extracted using the Voicebox toolbox, the binary regressor time-locked to the offset of each word in the audio (`word rate`), and the unigram frequency of each word (`frequency`), estimated using the Google ngrams and the SUBTLEX corpora for English (Brysbaert & New, 2009) and Chinese (Cai & Brysbaert, 2010). These regressors were convolved with SPM12’s (Penny et al., 2011) canonical HRF function and matched the scan numbers of each section. At the group level, the contrast image for English and Chinese were examined by a factorial design matrix. An 8 mm full-width at half-maximum (FWHM) Gaussian smoothing kernel was applied on the contrast images from the first-level analysis to counteract inter-subject anatomical variation. Statistical significance was determined by a cluster-based permutation test (Maris & Oostenveld, 2007), thresholded at *p* < .05 *FWE*.

A similar two-stage regression analysis was conducted to find the significant spatiotemporal clusters that are correlated with third person pronoun processing. We applied the same regression model to each participant’s single-trial source estimates at each timepoint of the whole 500 ms time window. This resulted in a *β* coefficient for each variable at each source and each timepoint for each subject. The source estimates were resampled to 100 Hz. At the second stage, we performed a one-sample *t*-test on the distribution of the *β* values for the binary third person pronoun regressor across subjects, again at each source and each timepoint. Clusters were then formed based on the *t*-values that were contiguously significant through time and space, at a level of *p* < .05. Only clusters that contained a minimum of 10 sources and spanned at least 10 ms were entered into a cluster-based spatiotemporal permutation test (Maris & Oostenveld, 2007). This involved randomly shuffling 0 and the *β* coefficient for each participant, repeating the mass univariate one-sample *t*-test and cluster formation within the 0-500 ms analysis window. This procedure was performed 10,000 times, resulting in a distribution of 10,000 cluster-level statistics. Each of the observed clusters was subsequently assigned a p-value based on the proportion of random partitions that resulted in a larger test statistic than the observed one. The GLM analysis was performed with MNE-python (v.0.19.2) (Gramfort & et al., 2014) and Eelbrain (v.0.25.2) (Brodbeck & et al., 2020).

### 2.8 RSA within the fROIs

For the fMRI data, the significant clusters from the GLM analyses were used as the fROIs to compare the five models’ relatedness to the brain data (see Figure 4c). For each subject in each group, we first extracted all the brain scans after the occurrence of each pronoun by aligning them with the offset of each pronoun in the audiobook. We added 5 seconds to the offset to capture the peak of the hemodynamic response function. The resulting English dataset contains 588 fMRI scans and the Chinese dataset contains 493 fMRI scans. We then regressed out the effects of intensity, f0, word rate and word frequency by subtracting from the brain data the four regressors multiplied by their beta values. Next, we calculated the RDMs between the brain activity patterns using 1 minus the Pearson’s correlation, computed across voxels.

**Figure 4:**
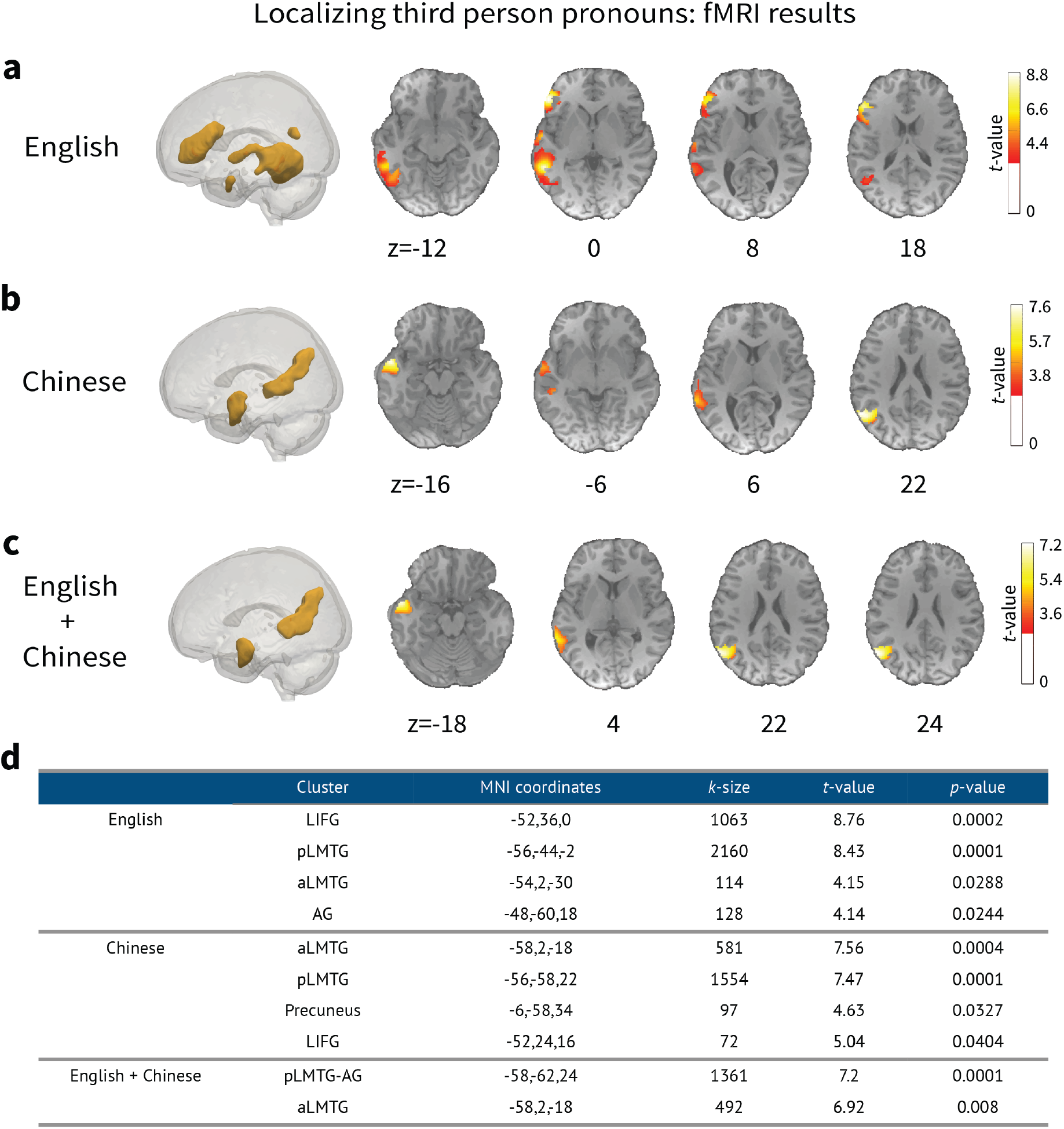
GLM results of the fMRI data for third person pronoun processing in English and Chinese. **a** Significant clusters of the English fMRI data. **b** Significant clusters of the Chinese fMRI data. **c** Significant clusters of the English and Chinese fMRI data. **d** MNI coordinates and their statistics, thresholded at p<.05 *FWE* with a cluster-based permutation test.

The complexity metrics derived from the Hobbs, Centering, ACT-R, ELMo-Coref and BERT-Coref models for each pronoun in the English and Chinese fMRI stimuli were used to construct the model RDMs, computed as the euclidean distance between each metric value. If multiple pronouns occur within one fMRI scan, the metrics were summed up. The English model RDMs are 588× 588 matrices and the Chinese model RDMs are 493× 493 matrices.

To compare the five models’ ability to explain the fMRI data RDM within the two fROIs, we calculated the Spearman’s rank correlation between the data RDMs and each model RDM for each subject in each group. Statistical significance was determined using a one-sided signed-rank test (Wilcoxon, 1945) across each subject’s correlation values. Pairwise comparisons between each model’s relatedness to the brain data were also tested using a one-sided signed-rank test. FDR correction was applied for multiple comparisons across fROIs and model pairs.

For the MEG data, we subset the source estimates of the pronouns within the lateral and medial clusters derived from the regression analyses for third person pronoun processing. At each timepoint, we computed the MEG data RDM for each pronoun as 1 minus Pearson’s correlation among the MEG data. The complexity metrics derived from the five models for each pronoun in the MEG stimuli were used to construct the model RDMs, computed as the euclidean distance between each metric value. The model RDMs are 51 ×51 matrices.

We calculated the Spearman’s rank correlation between the MEG data RDM and each model RDM at each timepoint for each subject. Statistical significance was determined using a cluster-based permutation *t*-test (Maris & Oostenveld, 2007) with 10,000 permutations across each subject’s correlation map. Clusters were formed based at a level of *p* < .05. Only clusters that contained a minimum of 10 sources and spanned at least 10 ms were entered into a cluster-based spatiotemporal permutation test. FDR correction was applied for multiple comparisons across fROIs and models.

## 3 Results

### 3.1 Localizing pronoun processing in space and time

Figure 4 summarizes the GLM for for the English and Chinese fMRI data. The English data showed significant clusters correlated with the occurrence of third person pronouns in the left inferior frontal gyrus (LIFG; *p* = 0.0002, *k* = 1063), the anterior LMTG (*p* = 0.00288, *k* = 114), the posterior LMTG (*p* = 0.0001, *k* = 2160) and the AG (*p* = 0.0244, *k* = 128). The Chinese data showed similar LIFG (*p* = 0.0404, *k* = 72) and aLMTG (*p* = 0.0004, *k* = 581) clusters and a cluster spanning the posterior LMTG and AG (*p* = 0.0001, *k* = 747). There is an additional Precuneus activation (*p* = 0.0327, *k* = 97) for the Chinese data (see Figure 4b). The significant regions common to both the English and Chinese data are the aLMTG and the pLMTG-AG clusters (*p* = 0.0001, *k* = 1361 and *p* = 0.008, *k* = 492, respectively). No significant cluster was found for the contrast between the English and Chinese groups.

The MEG data revealed one significant cluster of 401 sources for third person pronouns from 150 to 250 ms (*p* = 0.033) after the onset of the word. The cluster covered regions including the left posterior temporal cortex, the angular gyrus and the left medial parietal lobe (Figure 5a,b). Figure 5c,d show the timecourses of the mean activation for third person pronouns and all other words averaged over the lateral and the medial parts of the cluster. Compared to other words, third person pronouns elicited higher activity in the significant time window.

**Figure 5:**
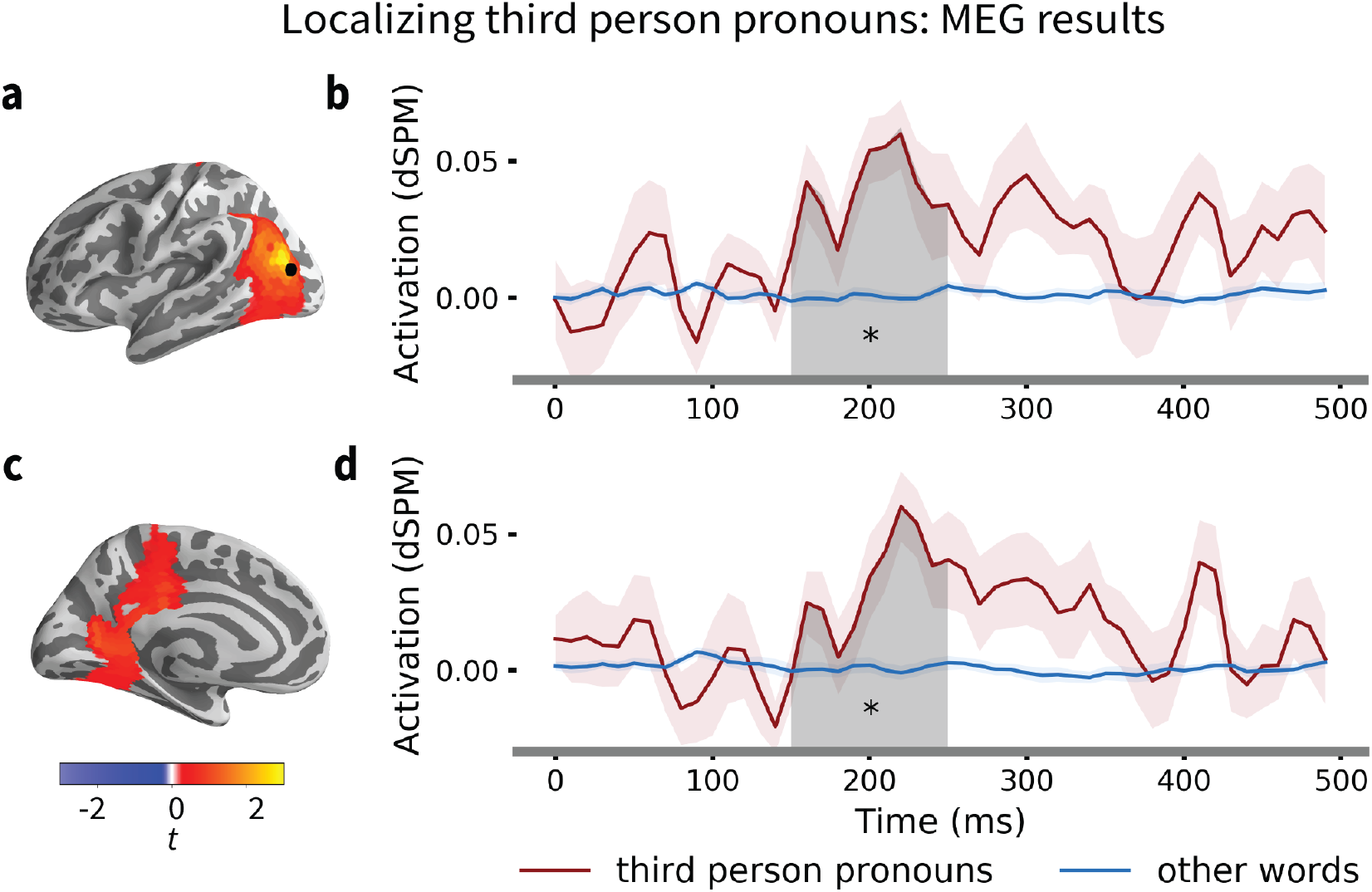
**a** Lateral part of the significant cluster for third person pronoun processing. **b** Timecourses of responses for third person pronoun and all other words averaged over the lateral cluster. **c** Medial part of the significant cluster for third person pronoun processing. **h** Timecourses of responses for third person pronouns and all other words averaged over the medial cluster. The shaded region indicates the significant time window from 150-250 ms after the word onset (p = 0.033).

Both our fMRI and MEG results showed significant LMTG activity, consistent with previous findings on the neural correlates of pronoun processing (Hammer et al., 2007, 2011; Miceli et al., 2002). Our MEG results showed additional activity in the left medial parietal lobe, which also replicated previous MEG results (Brodbeck et al., 2016; Brodbeck & Pylkkänen, 2017). We extracted the aLMTG and the pLMTG-AG clusters common to both the English and Chinese fMRI data as the functional regions of interests (fROIs) to compare computational models for pronoun resolution with the fMRI data using RSA. Similarly, we split the significant cluster from the MEG results into a lateral part and a medial part and used them as the fROIs to evaluate the models against the MEG data.

### 3.2 Comparing model predictions within the fROIs of the fMRI data

We performed RSA (Kriegeskorte et al., 2008) for each model within the fROIs for third person pronoun processing to compare the model predictions within the significant clusters for third person pronoun resolution in English and Chinese. RSA characterizes a representation in the brain or a computational model by the representational dissimilarity matrix (RDM) of the brain activity patterns or the model predictions. A model is tested by comparing the RDM it predicts to that of a measured brain region. We used the complexity metrics derived from the five models for each third person pronoun in the fMRI stimuli to construct the model RDMs. The data RDMs were computed as 1 minus Pearson’s correlation among the BOLD responses. Figure 6a,c show the RDMs for the Hobbs, Centering, ACT-R, ELMo-Coref and BERT-Coref model in English and Chinese. Spearman’s rank correlation among the five models is shown in Figure 6b,d.

**Figure 6:**
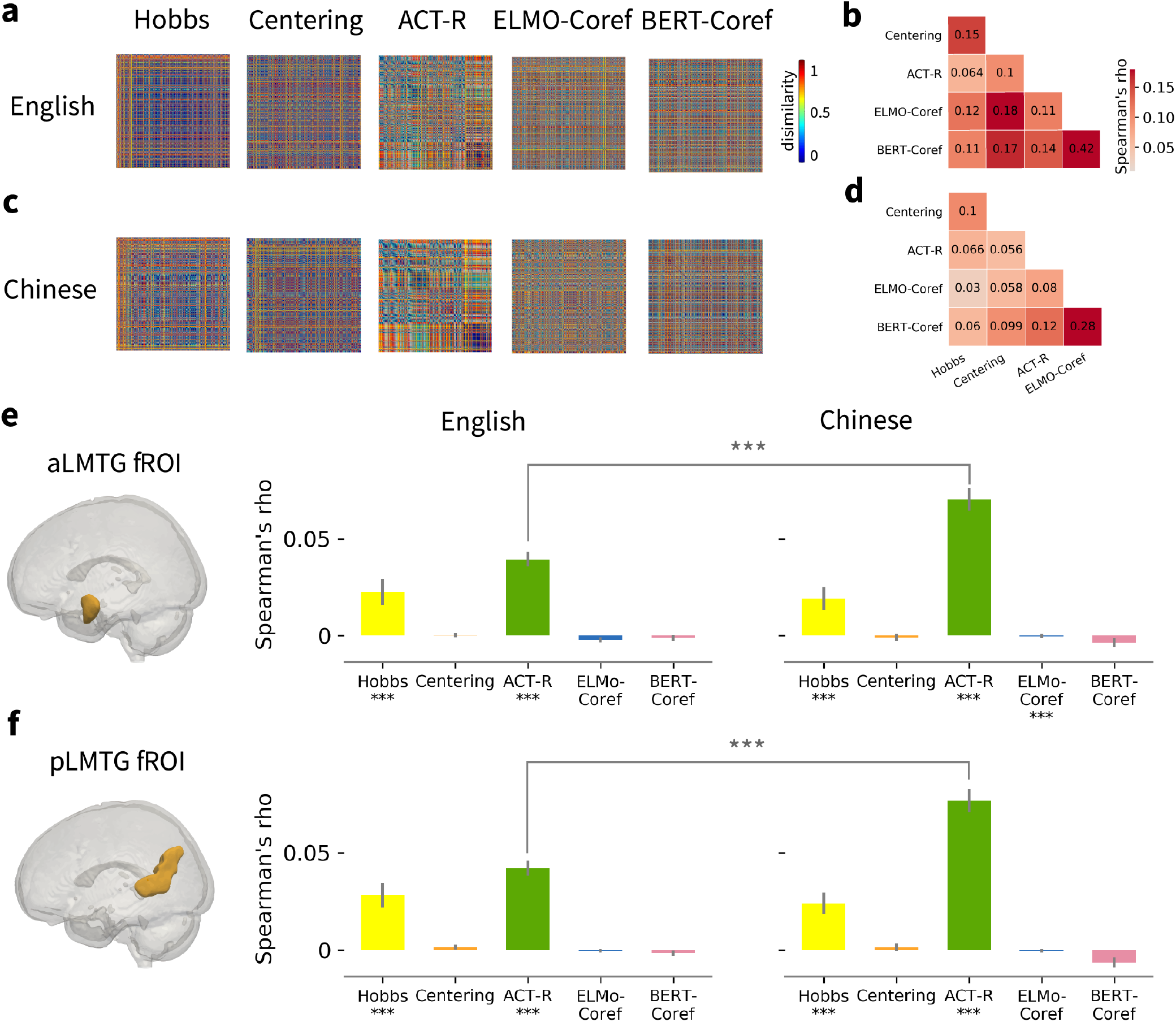
ROI-based RSA results for the fMRI data. **a** The RDMs for the Hobbs, Centering, ACT-R, ELMo-Coref and BERT-Coref model in English. Each RDM was separately rank-transformed and scaled into [0,1]. **b** Spearman’s rank correlation matrix for the five model RDMs in English. **c** The model RDMs in Chinese. **d** Spearman’s rank correlation matrix for the five model RDMs in Chinese. **e** Relatedness of the five model RDMs to the brain data RDM within the aLMTG fROI, averaged across subjects in each group. **f** Relatedness of the five model RDMs to the fMRI data RDM within the pLMTG-AG fROI, averaged across subjects in each group. * below the model names denotes significant correlation. * above the bars denotes significant group difference. FDR correction was applied for multiple comparisons across fROIs and models. *** *p* < .0001.

The RSA results for the English and Chinese data are shown in Figure 6e,f. For the English data, the ACT-R model showed the highest Spearman’s rank correlation with both the aLMTG and pLMTG-AG activities, averaged across subjects (*rho* = 0.039, *p* < .0001 and *rho* = 0.042, *p* < .0001, respectively). The Hobbs model was also significantly related to both fROI activities (*rho* = 0.023, *p* = 0.0015 and *rho* = 0.028, *p* < .0001, respectively). The Centering, ELMo-Coref and BERT-Coref model RDMs were not significantly correlated with either of the fROI activities (Centering: *rho* = 0.0004, *p* = 0.47 and *rho* = 0.0015, *p* = 0.23; ELMo-Coref: *rho* = −0.0021, *p* = 0.13 and *rho* = −0.0004, *p* = 0.57, BERT-Coref: *rho* = −0.0016, *p* = 0.87 and *rho* = −0.0016, *p* = 0.87).

For the Chinese data, the ACT-R model also had the highest mean correlation value with both the aLMTG and pLMTG-AG regions (*rho* = 0.07, *p* < .0001 and *rho* = 0.077, *p* < .0001, respectively). The Hobbs model was also significant for both fROIs with a *rho* value of 0.019 (*p* = .002) and 0.024 (*p* < .0001), respectively. The ELMo-Coref model showed a significant but very low correlation with the aLMTG activity (rho = 0.003, *p* = 0.005) but not the pLMTG-AG activity (*rho* = 0.0014, *p* = 0.66). The Centering and BERT-Coref models were not significant for either of the fROIs (Centering: *rho* = −0.001, *p* = 0.66 and *rho* = 0.0014, *p* = 0.7; BERT-Coref: *rho* = −0.0038, *p* = 0.94 and *rho* = −0.0064, *p* = 0.99).

A two-sample t-test between the English and Chinese speakers revealed a significantly higher correlation of the ACT-R models with the Chinese fMRI data in both the aLMTG and pLMTG-AG clusters (*t*(82) = 4.6, *p* < .0001 and *t*(82) = 5, *p* < .0001), suggesting a better fit of the ACT-R model for pronoun resolution in Chinese.

### 3.3 Comparing model predictions within the fROIs of the MEG data

For the MEG data, we subset the source estimates of the pronouns within the lateral and medial fROIs derived from the GLM analysis for third person pronoun processing. At each timepoint, we computed the MEG data RDM for each pronoun as 1 minus Pearson’s correlation between the MEG source estimates for all the third person pronouns. The complexity metrics derived from the five models for each pronoun in the MEG stimuli were used to construct the model RDMs, computed as the euclidean distance between each metric value. Figure 7a shows the five model RDMs, each separately rank-transformed and scaled into [0,1]. Figure 7b shows the Spearman’s rank correlation matrix for the five model RDMs.

**Figure 7:**
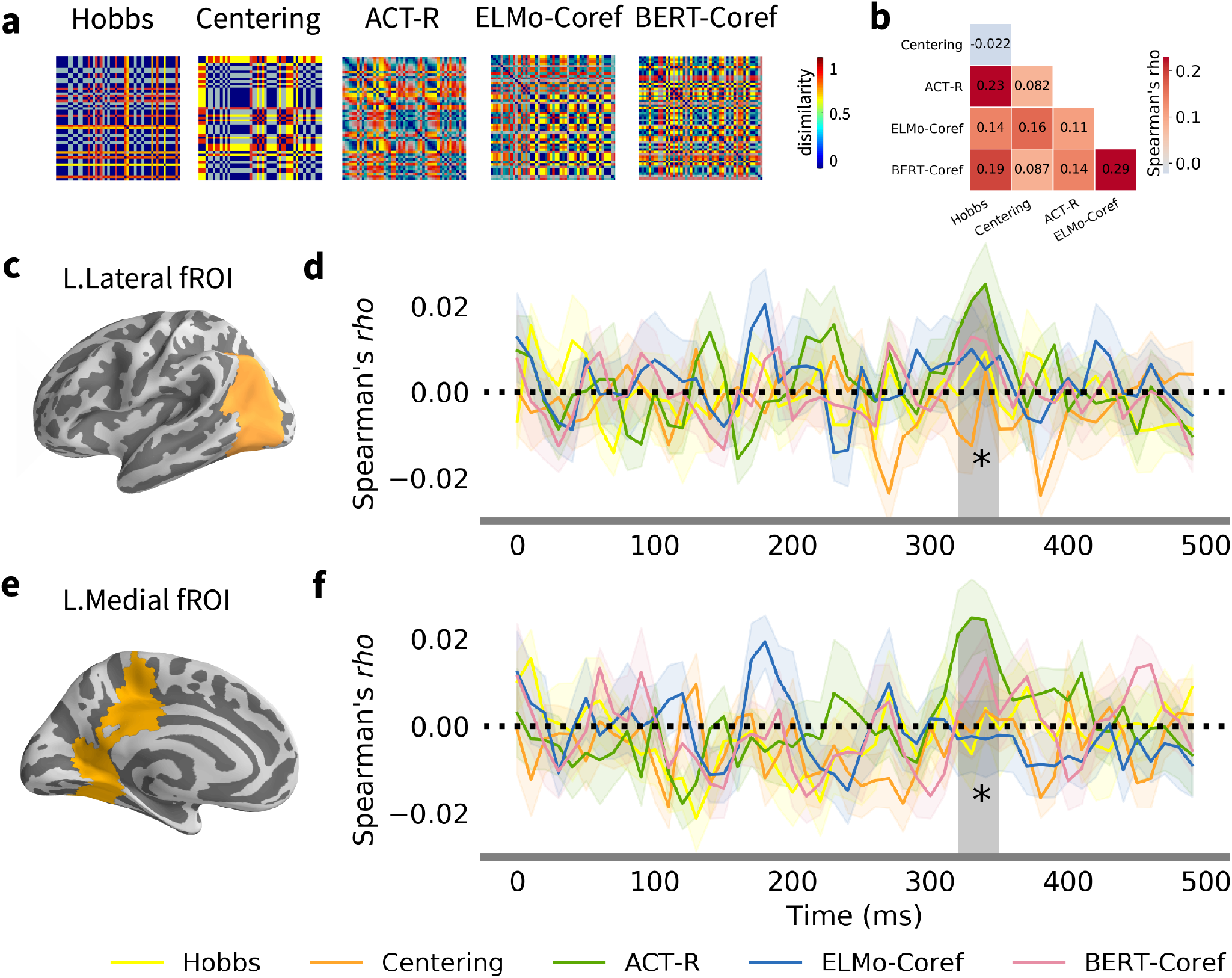
ROI-based RSA results for the MEG data. **a** The RDMs for the Hobbs, Centering, ACT-R, ELMo-Coref and BERT-Coref model metrics for the MEG stimuli. Each RDM was separately rank-transformed and scaled into [0,1]. **b** Spearman’s rank correlation matrix for the five model RDMs. **c** fROI in the left lateral lobe derived from the GLM analysis for third person pronoun processing. **d** Timecourse of the relatedness of the five model RDMs to the brain data RDM within the lateral fROI, averaged across subjects in each group. ACT-R model is significantly correlated with the MEG data pattern in the lateral fROI from 320-350 ms (*p* = 0.049). **e** fROI in the left medial wall. **f** Timecourse of the relatedness of the five model RDMs to the brain data RDM within the lateral fROI, averaged across subjects in each group. The ACT-R model is significantly correlated with the MEG data in the medial fROI from 320-350 ms (*p* = 0.038). Shaded region denotes significant temporal cluster. * *p* < .05.

The RSA results showed a significant cluster for the ACT-R model from around 320 to 350 ms after the onset of the pronouns for both the lateral fROI (*p* = 0.049) and the medial fROI (*p* = 0.038). None of the other three models was significantly correlated with the MEG data patterns within either fROI (see Figure 7c-f).

## 4 Discussion

The aim of the study is to investigate both the neural correlates and the cognitive mechanisms underlying referential processing during naturalistic language comprehension. For the neural correlates of reference processing, both our fMRI and MEG results converged on the LMTG and the AG across English and Chinese. The LMTG has been previously associated with biological and syntactic gender processing (Heim et al., 2002; Hammer et al., 2007, 2011; Miceli et al., 2002), and morphological inflection (Longe et al., 2007). For example, Hammer et al. (2007) showed that German sentences with congruent biological and syntactic gender were correlated with increased activation in the LMTG; Miceli et al. (2002) reported the LMTG activity when the subjects were asked whether a written noun has a masculine or a feminine gender. Longe et al. (2007) found greater LMTG activity for plurals and inflected verbs compared to stem words. Both gender and inflections are relevant for pronoun resolution in English since English pronouns distinguish gender and Case (i.e., “she” for subject and “her” for object). However, we found that the LMTG is also implicated for pronoun processing in Chinese, where pronouns do not have gender (“she”, “he” and “it” are all pronounced as “ta” in Chinese) or Case inflections. We therefore suggest that the LMTG supports the retrieval of reference in general, consistent with Indefrey & Levelt’s (2004) model on language production where the LMTG is the site for lemma selection and retrieval. Furthermore, lesions in the LMTG led to selective difficulty in accessing nouns (Lambon Ralph et al., 2000; Miceli et al., 1991), supporting the LMTG’s role in retrieving references.

The left AG has been previously implicated for referential processing involving backward anaphora (e.g., “Because he_*i*_ extinguished the flames, the fireman_*i*_ saved the resident that arrived later.”) (Matchin et al., 2014). Structurally, the AG adjoins the visual, spatial, auditory, and somatosensory regions. This makes it the best candidate for a high-level, supramodal integration area in the human brain (Geschwind, 1965). There is ample evidence for the AG’s role in multi-modal and multi-sensory associations/integration for attention, episodic and semantic memory, and sentence level comprehension (Bonner et al., 2013; Davis & Yee, 2019; Dronkers et al., 2004; Geschwind, 1965; Humphreys et al., 2021; Ramanan et al., 2018; Seghier, 2013; Tyler et al., 2011). Lesions in this region led to a variety of deficit in sentence comprehension (see e.g., Dronkers et al., 2004), such as more “role reversal errors” (i.e., who did what to whom) in sentence-picture matching tasks where they heard sentences either in the active voice (e.g. “The horse chases the boy”) or the passive voice (e.g., “The boy is chased by the horse”) (Tyler et al., 2011). The left AG has also been implicated in combinatorial conceptual/semantic processing (Binder et al., 2009; Branzi et al., 2021; Fletcher et al., 1995; Humphries et al., 2007; Homae et al., 2003; Price et al., 2015; Xu et al., 2005). Humphries et al. (2007), for example, observed the left AG activity for semantically congruent sentences compared to semantically incongruent sentences, semantically congruent word lists, random word lists and pseudo-word lists; (Price et al., 2015) found increased AG activity for words of higher combinatorial strength compared to unrelated words; Branzi et al. (2021), using Transcranial Magnetic Stimulation (TMS), found that the left AG is critical for integrating context-dependent information during naturalistic language comprehension. Binder et al.’s (2009) meta-analysis on semantic processing also suggested that the AG is at the top of a processing hierarchy underlying concept retrieval and conceptual integration, and is therefore essential for discourse-level comprehension.

The MEG data showed an additional recruitment of the left medial parietal region for third person pronoun processing, and the Chinese fMRI data also showed a significant cluster in the left Precuneus. This medial parietal activity has been previously reported when retrieving referentially ambiguous pronouns, as in “Ronald told Frank that he …” (Nieuwland et al., 2007), or when the sentence contains two referents, as in “Jeremy and Lucy did some work on the house next door.” (Boiteau et al., 2014). These findings all suggest that the left medial parietal region may be responsible for tracking multiple referents. Almor et al. (2007) further suggested that the parietal regions might be originally devoted to perceptual organization where it tracks multiple objects in space. Indeed, Brodbeck et al. (2016) used a visual world paradigm in MEG and found the medial parietal activity for successful reference resolution. Therefore, one interpretation of the medial parietal activity in the MEG data and the Chinese fMRI data might be that there are more ambiguous pronouns in the MEG stimuli. Moreover, due to the lack of gender and case information, Chinese speakers may encounter more difficulty in processing referentially ambiguous pronouns.

The MEG data also provide information on the temporal dynamics of the neural activity for pronoun processing. We show that pronoun processing starts from 150 ms till 250 ms after the onset of the pronoun, consistent with Hauk et al.’s (2012) finding that the brain starts retrieving lexical and semantic information within around 200 ms of word onset in the LMTG. We are aware of only two MEG studies on referential processing in the literature Brodbeck & Pylkkänen (2017); Brodbeck et al. (2016), and our time window is a bit earlier than the 350-500 ms window previously reported. However, the previous MEG studies used a visual word paradigm with multiple candidate referents, which might be more difficult to resolve than the pronouns in our naturalistic listening setting. Moreover, the previous analysis time window was restricted to 200-500 ms after the target stimuli, which is different from the 0-500 ms analysis time window in our study.

To further understand the fine details of the cognitive processes underlying pronoun resolution in the brain, we leveraged computational models for pronoun resolution. We tested three knowledge-based symbolic models and two data-driven neural network models for pronoun resolution against both fMRI and MEG data. Our results all favor the ACT-R model (van Rij et al., 2013): For both the English and Chinese fMRI data, the ACT-R model showed the highest correlation with the BOLD response patterns within the aLMTG and pLMTG-AG fROIs; for the MEG data, the ACT-R model is the only model that showed significant correlation with the source-localized MEG data within the left posterior temporal lobe and the left medial parietal fROIs. Our results therefore support the memory-based account for pronoun resolution (Nieuwland & van Berkum, 2006), which emphasizes the role of memory resources while retrieving the correct references.

Working memory capacity has long been associated with individual differences in measures of language comprehension in the psycholinguistic literature, and there is a wealth of data suggesting that individuals who perform better on a typical verbal working memory task like the Reading Span task also perform better on both off-line and on-line measures of language comprehension (e.g., Daneman & Carpenter, 1980, 1992). The concept of “memory load” has also been related to complexity in sentence processing. For example, Gibson (1998) claimed that object relative clauses such as “The reporter that the senator attacked disliked the editor.” are more difficult to process than subject relative clauses such as “The reporter that attacked the senator disliked the editor.”, because the syntactic structure of object relative clauses leads to higher memory load: The distance between the subject (“the reporter”) and its “trace” position is longer in object relative clauses than in subject relative clauses (i.e., the trace position for “the reporter” in the object relative clause is after the verb “attack” whereas the trace in the subject relative clause is before “attack”).

The memory-based account for pronoun resolution employ the a similar concept of memory resource in psycholinguistics, it views pronoun resolution as retrieving the most salient entities from declarative memory. Built using the primitives of the memory module in the cognitive architecture ACT-R (J. R. Anderson, 2005), the ACT-R model incorporates the frequency and recency effects of memory decay, with a boost of activation spread from previous mentions of the entities that occupy the subject position of a sentence. Although the Hobbs and the ELMo-Coref model also incorporate a notion of recency, and the Centering model contains the feature of subjecthood, the parameters for the ACT-R formula was directly developed using relevant fMRI data (J. R. Anderson, 2005). Our correlational results suggest that this algorithm indeed correlates better with the brain’s computation states than other non-brain-inspired models.

The two deep neural network models for coreference resolution did not correlate well with the brain data. Deep learning models have led to significant advances in many aspects of natural language processing, and there has been considerable enthusiasm to relate the model architectures with the neural architectures for vision (e.g., Bao et al., 2020; Cadena et al., 2019; Kietzmann et al., 2019) and language (e.g., A. Anderson et al., n.d.; Toneva et al., 2021; Schrimpf et al., 2021). However, many of the deep learning models are not intended to match the human cognitive process. The two neural coreference models here, for example, treated coreference resolution as a clustering task, which is intuitively quite different from how humans comprehend pronouns. Moreover, both the ELMo and BERT architectures are bidirectional with information available from the following discourse, which the human listeners do not have access to during naturalistic listening comprehension. Therefore, although the neural network models performed quite well on the coreference task, they did not match the brain’s mechanism of referential analysis.

To sum up, we show that referential processing in human language is primarily localized at the LMTG, and we provide the first evidence that, in natural stories, brain activity for pronoun processing is best explained by the declarative memory module in the ACT-R cognitive architecture. Accordingly, reference resolution in rich contextual settings can be explained via domain-general, neurobiological mechanisms of memory retrieval in the human brain. Our work strengthens connections between cognitive neuroscience, psycholinguistics and natural language processing. By testing computational models rigorously against neuroimaging data, we also hope to provide insights on designing machines that learn and think like humans. We call for a joint force of cognitive neuroscience and artificial intelligence to explicate the intricate details of the human mind.

## Acknowledgements

This work is supported by the National Science Foundation under Grant No.1607441 and Grant No.1923144. The Chinese data collection is supported by the Jeffrey Sean Lehman Fund for Scholarly Exchange with China at Cornell University. We thank the Jiangsu Key Laboratory of Language and Cognitive Neuroscience at Jiangsu Normal University for help with data collection. We thank Maria Julieta Guzman and Samantha Wray for segmenting the MEG audio stimuli. We thank the High Performance Computing resources at NYUAD for computing support.

## Author contributions

J.H. designed the fMRI study. J.L. and L.P. designed the MEG study. J.L., J.H. and W.L collected the English fMRI data. J.L. collected the Chinese fMRI data and the MEG data. J.L. and S.W analyzed data. All authors wrote the paper.

## Supplementary Information

**Supplementary Figure 1:**
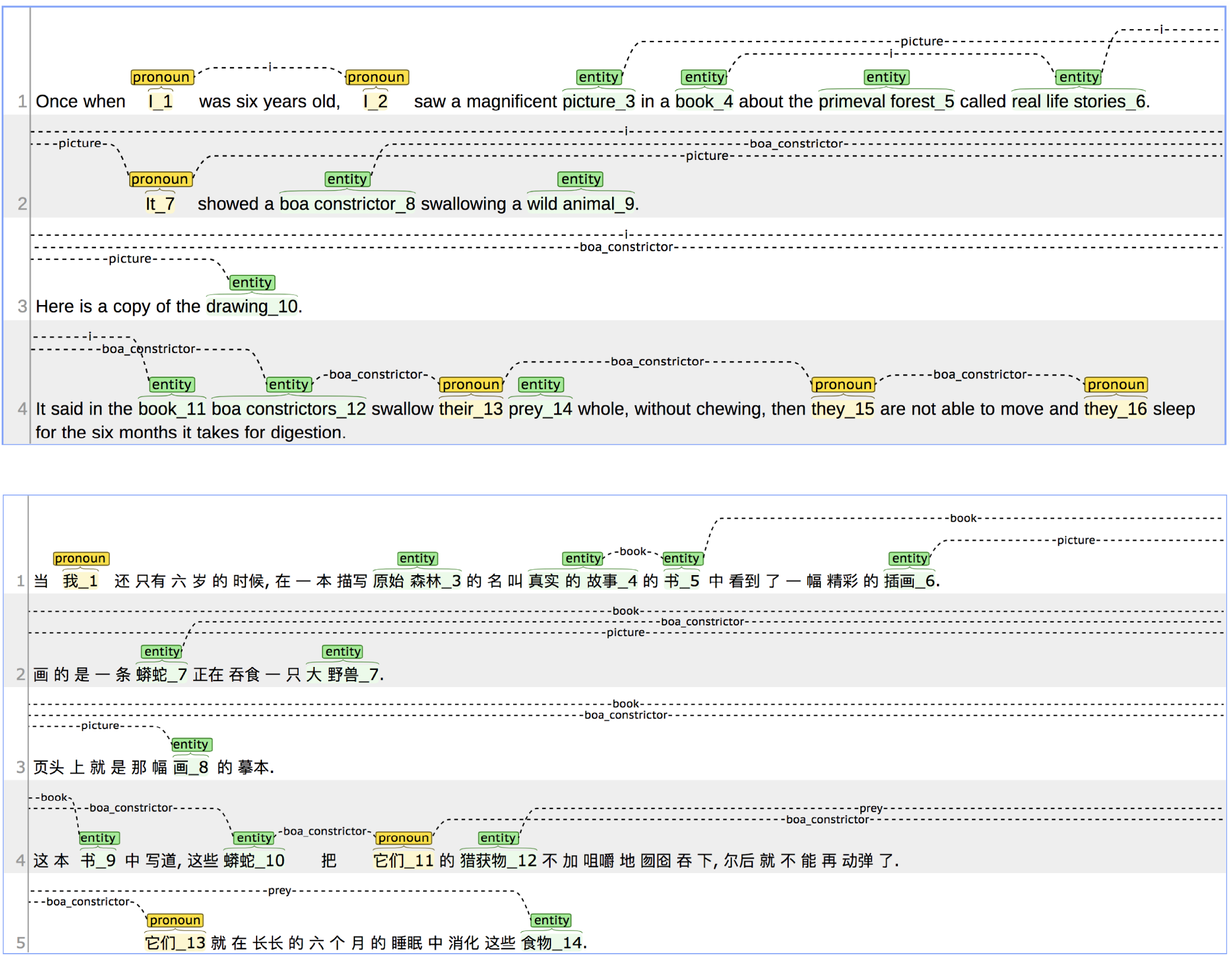
Sample annotations of pronouns and non-pronoun mentions in *The Little Prince* in English and Chinese, visualized using the annotation tool brat (Stenetorp et al., 2012).

**Supplementary Figure 2:**
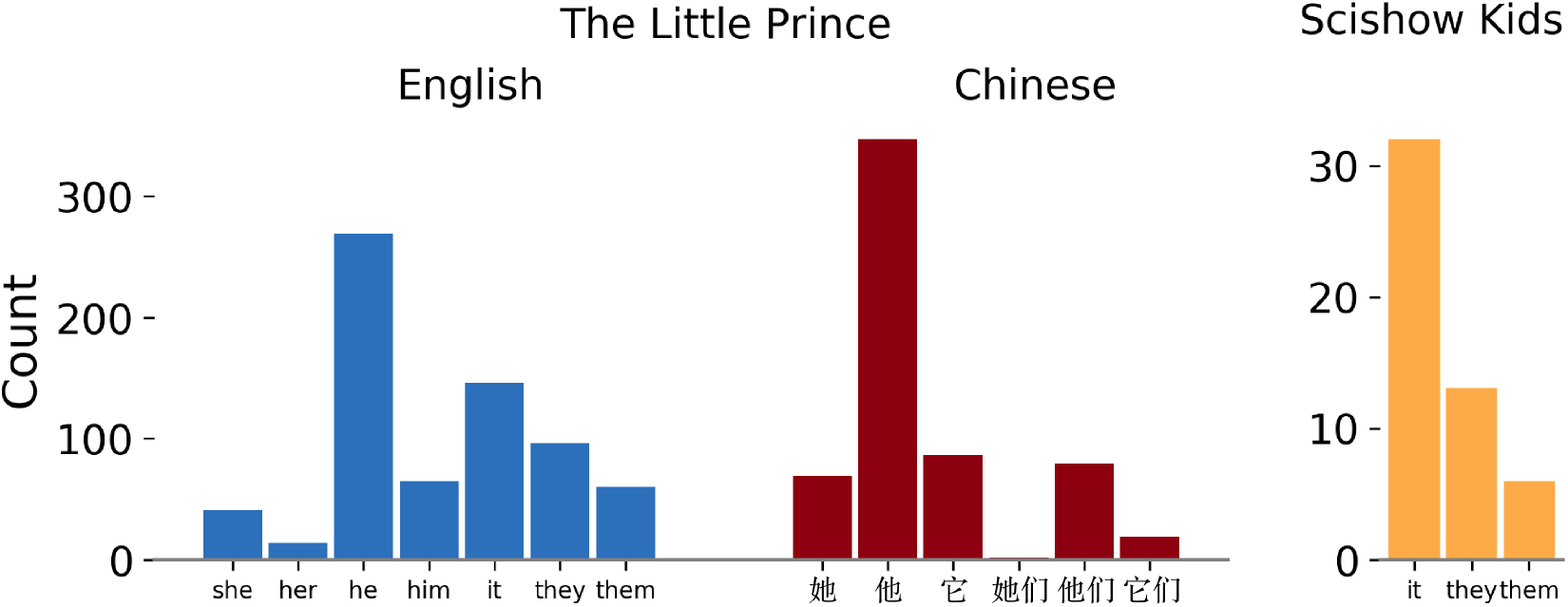
Counts for third person pronouns in the English and Chinese *The Little Prince and the SciShow Kids text*.

